# Rapid mode switching facilitates the growth of *Trichodesmium*: A model analysis

**DOI:** 10.1101/2023.07.14.549029

**Authors:** Meng Gao, Jamal Andrews, Gabrielle Armin, Subhendu Chakraborty, Jonathan P. Zehr, Keisuke Inomura

**Affiliations:** Graduate School of Oceanography, University of Rhode Island, Narragansett, Rhode Island, USA; Biological and Environmental Sciences Graduate Program, University of Rhode Island, Kingston, USA; Systems Ecology Group, Leibniz Centre for Tropical Marine Research (ZMT), Bremen, Germany; Department of Ocean Sciences, University of California, Santa Cruz, CA, USA

**Keywords:** *Trichodesmium*, nitrogenase, N_2_ fixation, photosynthesis, model, state switch, growth rate, carbon, nitrogen, O_2_ concentration

## Abstract

*Trichodesmium* is one of the dominant dinitrogen(N_2_)-fixers in the ocean, influencing global carbon and nitrogen cycles through its biochemical reactions. Although the photosynthetic activity of *Trichodesmium* fluctuates rapidly at the cellular level, the physiological or ecological advantage of this characteristic is not clear. Here we develop a metabolic model of *Trichodesmium* that can perform dinitrogen (N_2_) fixation during the daytime. We examined (1) the effect of the duration of switches between photosynthetic and non-photosynthetic cellular states and (2) the effect of the presence and absence of N_2_ fixation in photosynthetic states. Our results show that a rapid switch between photosynthetic and non-photosynthetic states increases *Trichodesmium* growth rates by improving metabolic efficiencies due to optimized C and N metabolism. Our results show the possibility and advantage of the rapid switch, providing a strategy for previous observations that all *Trichodesmium* cells can contain nitrogenase, which was previously considered to be a paradox. This study reveals the importance of fluctuating photosynthetic activity and provides a mechanism for daytime N_2_ fixation that allows *Trichodesmium* to fix N_2_ aerobically in the global ocean.

## Introduction

*Trichodesmium* is a cyanobacterial genus whose species have a multicellular filamentous morphology^1^ and are widely distributed in tropical and subtropical areas ^2–7^. It is also an important dinitrogen (N_2_)-fixing cyanobacterial genus in the global ocean, accounting for half of the N_2_ fixation in marine systems ^8,9^. In addition to fixing N_2_, *Trichodesmium* is photosynthetic and evolves oxygen (O_2_) ^10,11^. *Trichodesmium* plays an important role in ocean biogeochemical cycles ^12^ since it contributes to carbon (C) and nitrogen (N) cycling. Due to its ecological importance, many lab ^13–15^ and modeling ^3^ studies have been performed to understand its physiology, including N_2_ fixation strategies ^4,16,17^.

The N_2_-fixing enzyme (nitrogenase) with metal cofactors can be inactivated by trace levels presence of O_2 18,19_. Since cyanobacteria evolve O_2_ through oxygenic photosynthesis, they have to use physiological or morphological strategies to avoid the inactivation of nitrogenase ^16,18,20^. Several strategies in cyanobacteria have been reported ^18,20^, for example, the formation of specialized N_2_ fixation cells that lack oxygenic photosynthetic activity (heterocysts) ^16,21–23^. However, *Trichodesmium* does not form heterocysts^10,24^. Although a recent study found that it can fix N_2_ in the dark^25^, lots of previous studies suggest it appears to fix N_2_ and photosynthesize in the same cells during the daytime with light ^26–29^. It is still unresolved how *Trichodesmium* fixes N_2_ aerobically, in the light while evolving photosynthetic O_2_.

Although *Trichodesmium*’s N_2_ fixation mechanism is still unclear, results of previous studies suggested that photosynthetic activities can be regulated during cellular-level photosystem state transitions, which are on the order of 1 minute ^3,16,30,31^. Based on this photosynthesis 1-minute on/off switch, we developed two hypotheses: (H1) nitrogen fixation only occurs during a non-photosynthetic state, or (H2) trichodesmium may continue fixing N_2_ during photosynthesis.

As for H1, a model by Inomura et al. 2019 ^3^ suggested that the intracellular O_2_ may be decreased rapidly on the time scale of seconds without photosynthesis, which provides an opportunity for N_2_ fixation. The rapid recovery of nitrogenase can be supported by evidence from studies in other species, which have a protein (Shethna Protein II, FeSII) for conformational protection of nitrogenase ^32^. This protein can quickly respond to O_2_ and make the nitrogenase in an inactive but oxygen-tolerant state until recovery. However, such a protein (or its genes) has not been found in *Trichodesmium*, and reactivation of nitrogenase likely requires a much longer time than seconds to recover from inhibition ^33,34^. This argument leads to Hypothesis 2 (H2) that *Trichodesmium* may continue fixing N_2_ during photosynthesis. Studies show that under ambient oxygen concentrations, it takes more than a few minutes to deactivate nitrogenase ^32,34^. Thus, *Trichodesmium* might tolerate short time intervals at high O_2_ concentrations, for example, one minute. It takes tens of minutes to resynthesize nitrogenase ^35,36^, which excludes the possibility that cells are constantly resynthesizing nitrogenase as it is damaged, and repair cannot keep pace with inactivation.

Based on these hypotheses, we built a metabolic model of *Trichodesmium* (Figure 1) and ran it under two situations (H1 and H2) and two switch modes (1-minute and 6-hour) to answer three main questions: 1) How do the cells fix N_2_ even though they are producing O_2_? 2) How can mode switching influence growth rate and why? 3) Based on element (C and N) allocation, which switch mode is more efficient? The following model illustrates the mechanism of *Trichodesmium* N_2_ fixation and the associated advantages of the mechanism.

**Figure 1.**
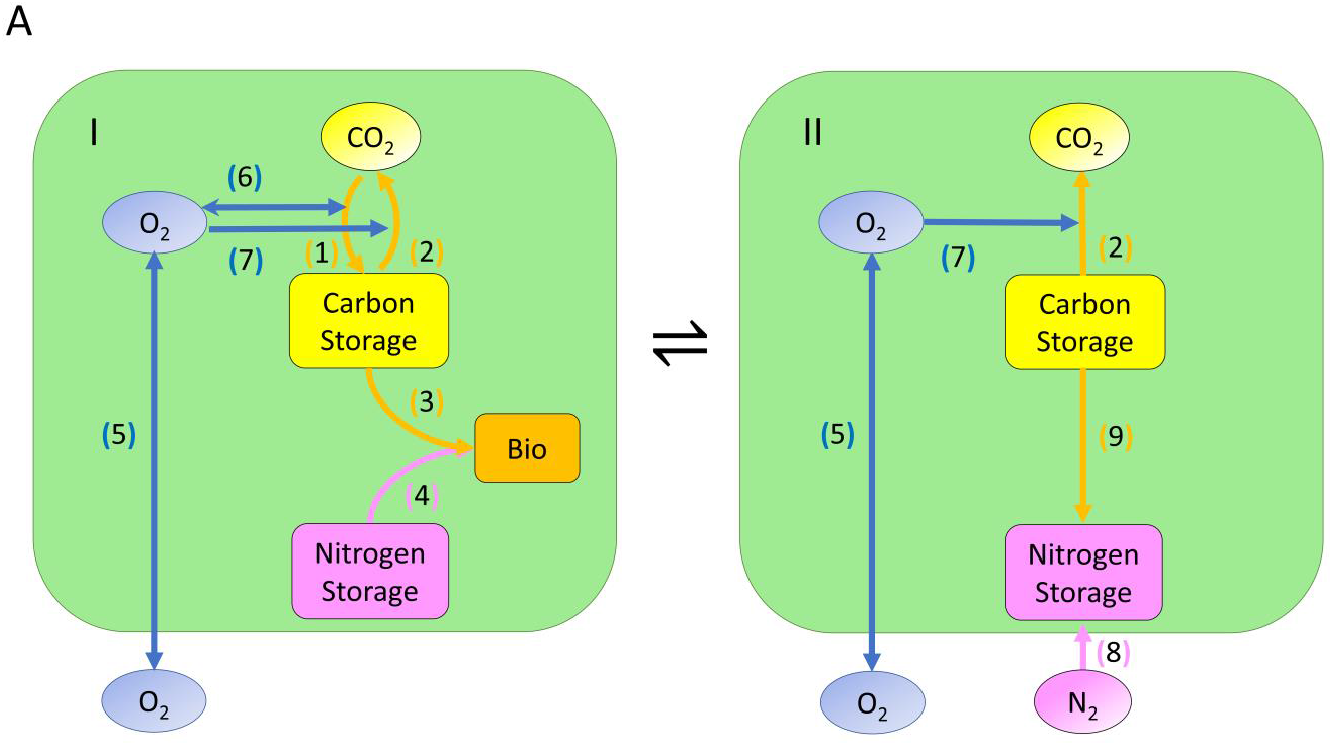
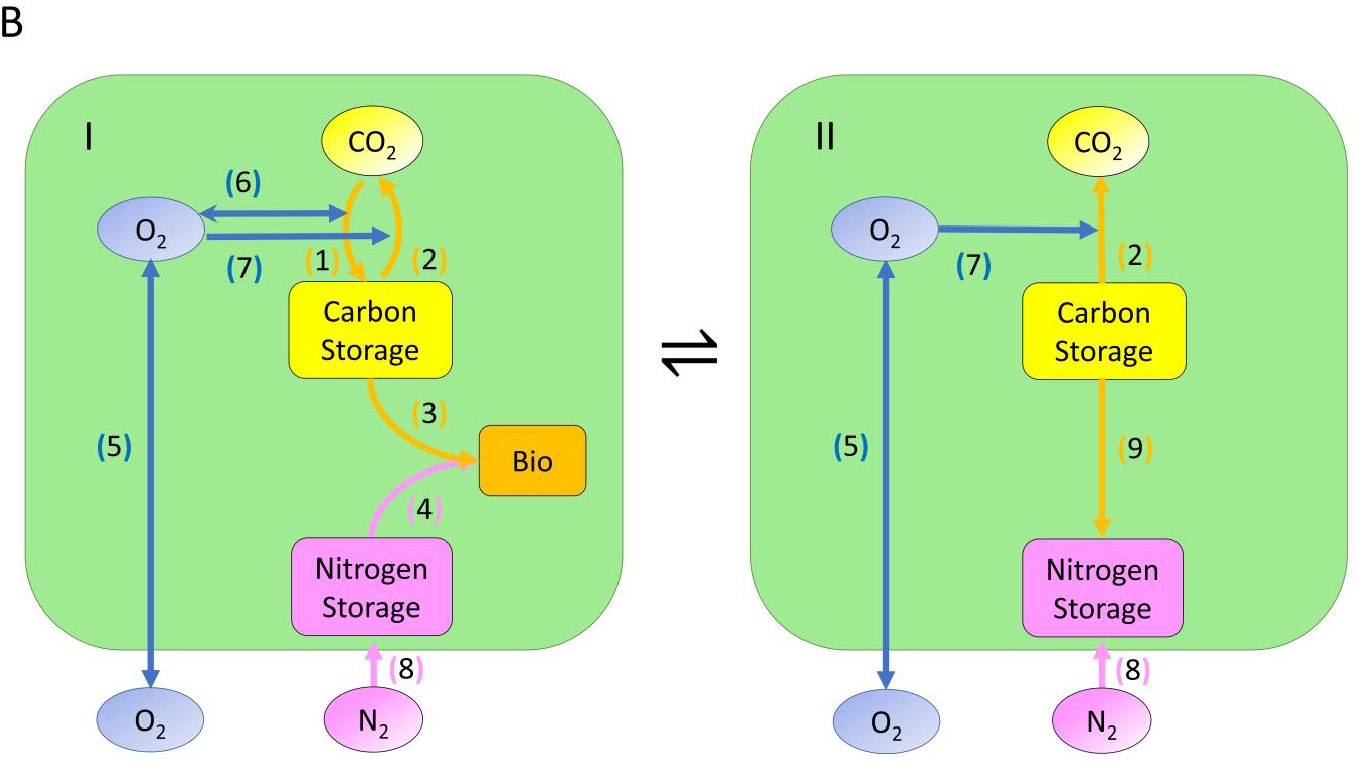
Schematic depiction of molecular pools and fluxes in the model. A. H1: Photosynthesis and N_2_ fixation occur at different times. B. H2: Photosynthesis and N_2_ fixation occur simultaneously. (I) Photosynthetic state. (II) non-photosynthetic (N_2_ fixation) state. The symbol ⇌ represents state transition. Pathways: (1) C fixation. (2) Respiration. (3) Consumption of C in growth. (4) Consumption of N in growth. (5) O_2_ diffusivity. (6) O_2_ changes in carbon fixation. (7) O_2_ changes in respiration. (8) N_2_ fixation. (9) Consumption of C for N_2_ fixation (for energy and electron). Different colors represent different element pools and fluxes, yellow is for C, pink is for N and blue is for O_2._ According to the real condition of *Trichodesmium* ^31^, we switched these two states every minute and every 6 hours. We ran the models for 12 hours. And we assumed that there is no growth in the N_2_ fixation state.

## Results and Discussion

### Overview of the model

We developed a physiological model of a *Trichodesmium* cell that performs photosynthesis to obtain carbon (C) and fix N_2_ to obtain nitrogen (N). The cell produces O_2_ during photosynthesis and uses both C and O_2_ to maintain respiration. Based on previous experimental evidence ^31^, here we assume that the cell can switch between photosynthetic and non-photosynthetic states. Accordingly, we built and examined two hypotheses H1 and H2; H1: N_2_ fixation happens only during N_2_ fixation state (Figure 1A, Video 1), whereas H2: N_2_ fixation happens in both N_2_ fixation state and photosynthetic states (Figure 1B, Video 2). Moreover, we considered two different switching modes based on the time duration of each state: a rapid mode, where the switching between two states happens every minute, and a slow mode, where the switching happens every six hours. To calculate cellular element dynamics (C, N, O_2_), we resolved molecular transport, photosynthesis, respiration, biosynthesis, and N_2_ fixation as the critical pathways. The following are the most important results.

### O_2_ fluctuation analysis

Our results show that O_2_ concentrations change dramatically as the state switches in both hypotheses. Figures 2A, B show that it took less than 1 second during the daytime to reach a steady O_2_ concentration after each switch. In the photosynthetic state, the O_2_ reached a high level, while in the N_2_ fixation state, O_2_ concentration decreased and remained low. The rapid decrease of O_2_ in the N_2_ fixation state might provide conditions to allow nitrogenase activity. Nitrogenase might (H1) get reactivated during low O_2_ conditions or (H2) tolerate the high O_2_ level for a short time during the beginning of the photosynthetic state. Based on these rapid O_2_ changes, *Trichodesmium* can photosynthesize and fix nitrogen (Figure S2) in the same cell, even with the temporal separation of only 1 minute.

**Figure 2.**
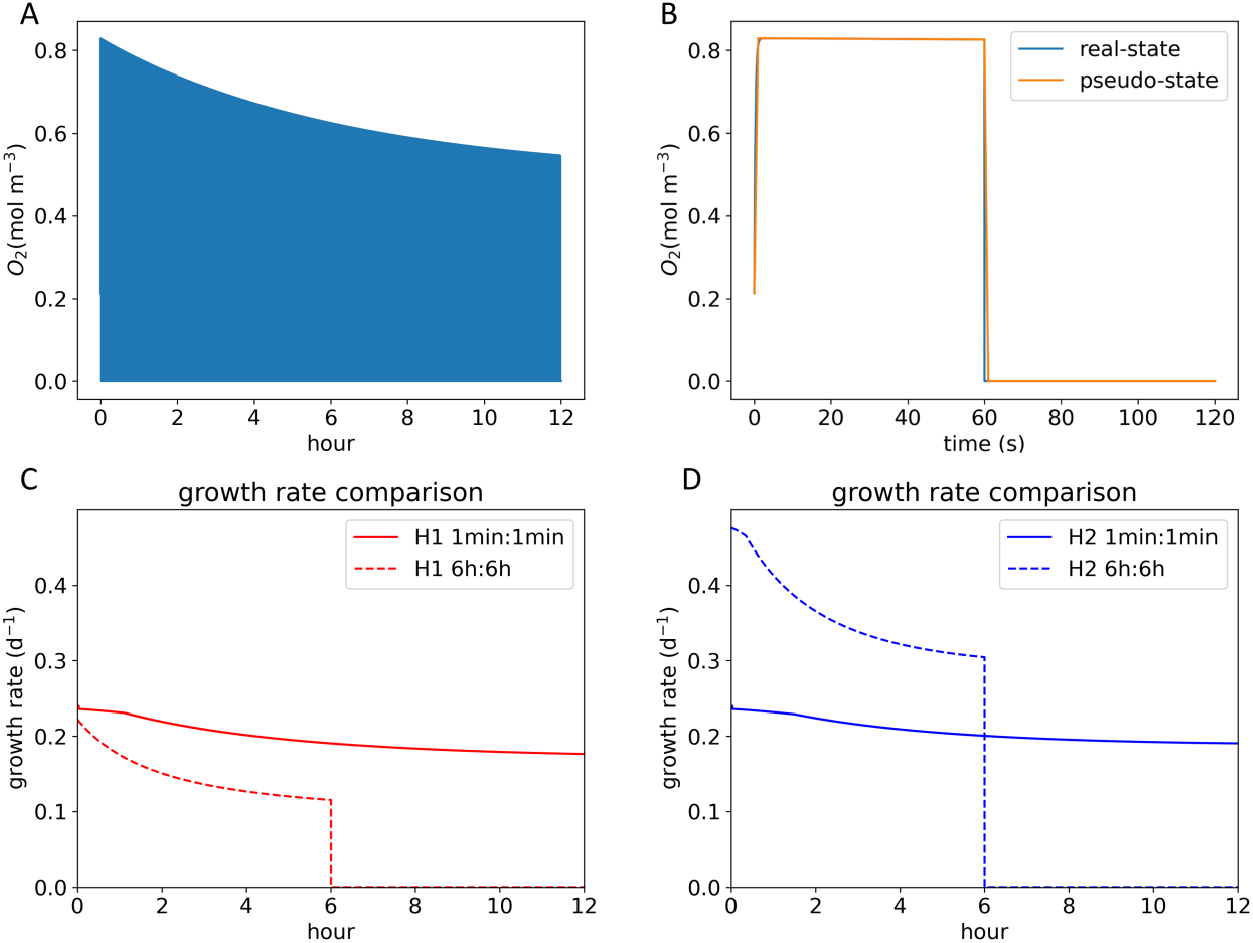
O_2_ level and growth rate changes in 12 hours. (A). Changes in O_2_ concentration in 12 hours (H2, and H1 pattern is similar in supplementary, Figure S1). (B). Changes in O_2_ concentration in 2 minutes (H2, and H1 pattern is similar in supplementary, Figure S1); 60 s for photosynthetic state and 60 s for non-photosynthetic state. Pseudo-state means that we used steady state conditions (O_2_ values do not change) to do the simulation. (C). Growth rate comparison of different modes in H1. (D.) Growth rate comparison of different modes in H2. For the rapid switch mode, we plot the growth rate by taking an average of 2 minutes.

Our simulated quick changes in O_2_ concentrations are similar to those of a previous modeling study ^3^, which estimated an extremely short residence time of O_2_ (the time to consume all O_2_ by respiration) in *Trichodesmium* cells on the order of 1 second. The rapid decrease results from the high respiration rate, as suggested by previous studies ^3,24,37,38^. Since N_2_ fixation requires substantial energy in the form of ATP (16 ATP per N_2_ fixed), it needs to be coupled with high aerobic respiration rates to provide ATP. The high aerobic respiration rates could explain our results that intracellular O_2_ level changes rapidly, and the O_2_ concentration is low throughout the N_2_ fixation state. In *Trichodesmium*, increased respiration in a cell might also reduce the plastoquinone pool and transmit negative signals to photosystem II (PSII), which would decrease photosynthesis and consequently the production of O_2_ ^10,37^. In addition to respiration, previous studies of N_2_ fixers suggested that lower O_2_ levels can also be maintained in the cell ^38,39^ by lowering O_2_ diffusivity ^3^ with the use of multiple membrane layers (gram-negative bacterium) ^40^ and extracellular polymeric substances (EPS) ^41–45^ (*Azotobacter vinelandii* and *Trichodesmium*), as well as using an alternative electronic transfer (AET) pathway (*Trichodesmium*)^37^.

### Growth rate comparison

We compared growth rates in 12 hours of daytime (with light) between rapid mode switching and slow mode switching (Figures 2C, D). For both H1 and H2, the average growth rates over 12 hours for rapid mode (H1: 0.20 d^-1^, H2: 0.21 d^-1^) are higher than those of slow mode (H1: 0.07 d^-1^, H2: 0.18 d^-1^). Figure 2C shows that under H1 the growth rate for the rapid mode is always higher than the slow mode, while Figure 2D shows that under H2 the first six-hour growth for the rapid mode is lower, but the remaining six-hour growth rate is higher. The difference between these two modes suggests that the rapid switch is a better strategy for *Trichodesmium*, which results from a better element supply and distribution inside the cell. To determine the reason for the higher growth rate, we simulated the element fate.

### Element fate and comparison

Here we compared C and N fates for two switching modes (Figure 3). During the rapid mode, under H1, the cell used a larger percentage of C in growth and storage and less in N_2_ fixation and respiration (Figure 3A). As for H2, a larger C percentage is used in growth and less in other pathways (Figure 3B). In terms of absolute values (indicated in x-labels), the total amount of C invested in cellular processes is higher in the rapid mode irrespective of H1 and H2. As for the N fate, results for both hypotheses showed that the rapid mode utilized more N in growth, in terms of both percentage and absolute values, than N storage (Figures 3C, D). The total utilization of N for the two modes is slightly different (H1: 0.041 and 0.042, H2: 0.084 and 0.081, unit: mol C mol N^-1^ d^-1^).

**Figure 3.**
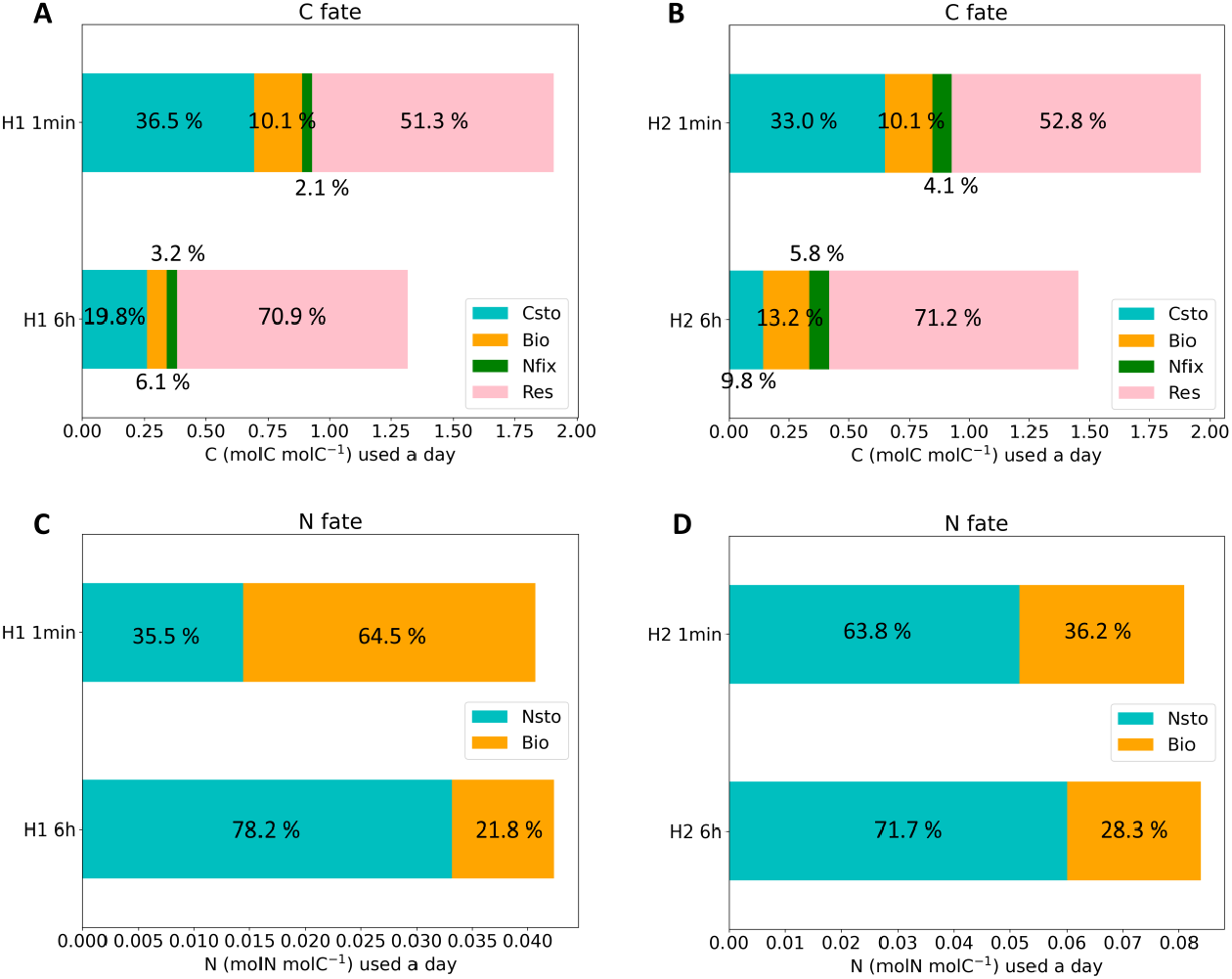
Element fate. C fate (C usage distribution in 12 hours) comparison for H1 (A) and H2 (B). N fate (N usage distribution in 12 hours) comparison for H1 (C) and H2 (D). Csto means C storage, Bio means element used in growth, Nfix means C used in N_2_ fixation, Res means C used in respiration, and Nsto means N storage, values in figures mean percentage values and stack lengths mean the absolute values.

We can explain our growth rate results based on the element comparison. To maintain the higher growth rate during the rapid mode switching, more C and N are used for growth (biomass) and less C for respiration. This shows that the rapid mode switching is a more efficient strategy with faster growth and less C lost, and thus *Trichodesmium* evolved the rapid mode switching to optimize benefits from photosynthesis and N_2_ fixation.

In summary, *Trichodesmium* switches every minute between photosynthesis and N_2_ fixation to achieve temporal segregation. This can happen because the O_2_ levels change so fast that nitrogenase can be protected during the N_2_ fixation state. The rapid mode change increases the growth rate by optimizing the distribution of C and N usage in cells and makes *Trichodesmium* a very successful N_2_ fixer and primary producer in the ocean and dominates the phytoplankton community in some areas ^17,46–49^.

### Comparison to previous studies and implications for future work

Previous research suggested that both spatial and temporal segregation mechanisms were used in *Trichodesmium*. Studies reported that *Trichodesmium* could form specialized cells (termed diazocytes) where N_2_ fixation was localized ^50^. However, it is still controversial since several studies reported that nitrogenase is randomly distributed in *Trichodesmium* cells or even in all cells, and a modeling study found that spatial separation is unnecessary ^37^. As a result, temporal separation is thought to be more necessary. Rapid state transition switches were shown in cellular-level fluorescence kinetics experiments ^30,31^, and our study reconciles these observations and provides possible mechanisms that facilitate daytime N_2_ fixation. Our study not only elucidates *Trichodesmium*’s O_2_ protection mechanisms but also reveals a strategy for non-heterocyst forming and daytime N_2_ fixing cyanobacteria, which can be a model for mechanisms in other species in addition to *Trichodesmium*. This model may provide a metabolic module of *Trichodesmium* for existing ecological models with *Trichodesmium* ^51–53^ for predicting their physiological response to the environment and the consequent ecological and biogeochemical impacts.

In our results, rapid mode switching results in higher efficiency in terms of resource use with more growth and less respiration compared to slow mode switching. Our simulated growth rates are consistent with experimentally observed *Trichodesmium* growth rates (0.1-0.5 d^-1^) in previous studies ^3,54,55^. Interestingly, nitrogenase can also be regulated by several environmental factors, and thus environmental factors may be important in affecting *Trichodesmium* N_2_ fixation and growth in the ocean. For example, ocean acidification ^56,57^ and nutrients like iron ^56,58^ have been found to influence nitrogenase efficiency. Future research on how environmental factors affect nitrogenase and state transitions will be important for predicting the response of *Trichodesmium* to climate change.

## Conclusion

With a mechanistic model of *Trichodesmium*, we found that rapid mode switching could facilitate N_2_ fixation because cellular O_2_ concentrations can decrease to a very low level in 1 second. This switch mode can also explain why most *Trichodesmium* cells contain nitrogenase and how they can fix N_2_ during the day. The rapid mode switching also keeps C and N concentrations flexible in cells to optimize C and N allocation, thus facilitating the growth of *Trichodesmium*. Our results show that switching rapidly is an efficient strategy for non-heterocyst-forming cyanobacteria and daytime N_2_-fixers, which can be a reference for further study in ocean N_2_-fixers.

## Materials and Methods

Here, we describe the ecological model, which includes the photosynthetic state and non-photosynthetic (N_2_ fixation) state of marine phytoplankton *Trichodesmium* (Figure 1). We calculate cellular C, N, and O_2_ concentrations during the half of the diurnal cycle using diffusivity, photosynthesis, respiration, and biosynthesis. Based on our two hypotheses, photosynthesis and N_2_ fixation can happen at the same time in H2, whereas they are separated by time in H1. In the following, first, we describe photosynthetic states for the two hypotheses and then the N_2_ fixation states. The following are the equations we used to build the model.

### Photosynthetic state

*H1*. To model phytoplankton C changing rate 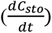 (Equation (1)) in the photosynthetic state, we assumed that it could be described as the difference between the C fixing rate (*F*_*cfix*_) and C consuming rate. This consumption includes respiration (*F*_*Bio*_E) and growth (*F*_*Bio*_), where E is the ratio of biomass to respiration production.

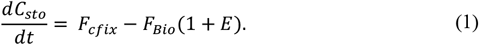

To estimate cellular O_2_ flux 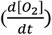, we included diffusivity, photosynthesis, and respiration pathways (Equation (2)). We used a product of diffusivity coefficient (A) and O_2_difference ([*O*_2_]_*E*_ - [*O*_2_]) to represent the O_2_ change in diffusivity between the extracellular (E) and intracellular environment. 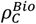 and 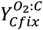 are cell carbon density and O_2_ to C ratio. Here we used them to quantify the O_2_ change in carbon fixation (*F*_*cfix*_) and respiration (*F*_*Bio*_*E*).

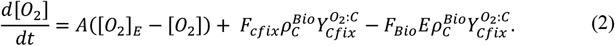

Next, we calculated N flux 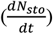 (Equation (3)) which only changes due to the consumption of nitrogen in growth in the photosynthetic state of *Trichodesmium*. Here, we used growth (*F*_*Bio*_) times the N to C ratio in cells 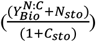. In this term, 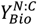 represents N to C ratio in biomass. We can also describe it as the N concentration when we use mol N mol C^-1^ as the unit of N content. We added the amount of N in the storage, *N*_*sto*_, to it to calculate the whole N concentration 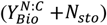. Here we used (1 + *C*_*sto*_) to represent the total C concentration in the cell. 1 here means the original functional C in cells and *C*_*sto*_ means C storage in cells.

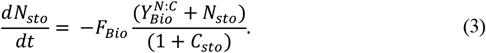

In Equation (4), we assumed that C fixation rate (*F*_*cfix*_) could increase with light intensity (*I*) but becomes saturated when it reaches a maximum 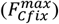. Here, *A*_*i*_ represents the light saturation coefficient.

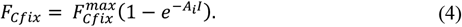

In Equation (5), we considered C and N cellular concentrations as two factors limiting biomass production. Due to Liebig’s law of minimum, growth is determined by the scarcest factors. Besides, biomass production can increase with C and N concentrations but will reach saturation at a high value. As a result, the form of the equation resembles Monod kinetics. Therefore, we calculated the actual biomass production, *F*_*Bio*_, as the minimum of available 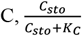, and 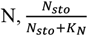, for biosynthesis. Here, *K*_*C*_ means half-saturation concentration for C and *K*_*N*_ means half-saturation concentration for N.

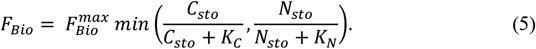

*H2*. Here N_2_ fixation happens during photosynthesis and thus we modified Equations (1) and (2) accordingly:

To calculate C changing rate, we added N_2_ fixation as another source of expenditure of C (Equation (6)). We included two more terms: 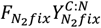 and 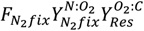, which respectively mean the C usage in N_2_ fixation and the C usage in respiration of N_2_ fixation. Here, 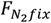 means N fixation rate, 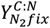 represents C to N ratio in N_2_ fixation, 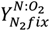 represents N to O_2_ ratio in N_2_ fixation, and 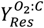 means O_2_ to C ratio in respiration.

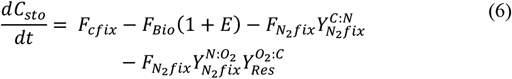

To revise the O_2_ flux equation, we subtracted a term 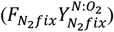 representing the respiratory cost of O_2_ during N_2_ fixation (Equation (7)).

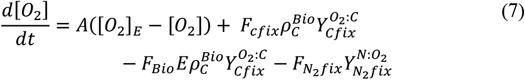

### Non-photosynthetic (N_2_ fixation) state

This state is the same for both the hypotheses H1 and H2. To calculate C changing rate in N_2_ fixation state, we assumed that it could be affected by two pathways: N_2_ fixation 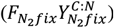 and respiration 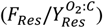. Here, 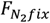 and *F*_*Res*_ represent N_2_ fixation rate and respiration rate, respectively. And we use C to N ratio 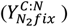 in N_2_ fixation and O_2_ to C ratio 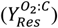 in respiration to transform the units.

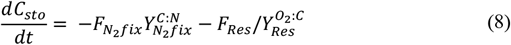

We calculate respiration rate (*F*_*Res*_) in Equation (9) by assuming that it increases with cellular O_2_ concentration ([*O*_2_]) and can reach a maximum rate 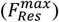.

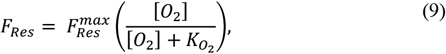

where, 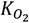 means half-saturation concentration of O_2_.

The only pathway affecting N changing rate 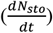 is N_2_ fixation 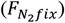, which is calculated in Equation 10. N_2_ fixation depends on cellular C storage (*C*_*sto*_). We assume that N_2_ fixation increases with increasing C storage and saturates due to physiological constraints to the maximum limit of N_2_ fixation rate 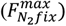 as:

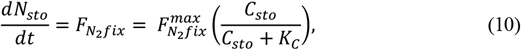

where *K*_*C*_ is half-saturation constant of C storage.

The cellular O_2_ flux is obtained by subtracting the respiratory consumption from the diffusive O_2_ input:

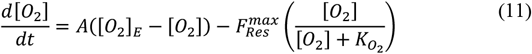

We calculated all of the element dynamics in the two states by simplification of the Taylor Expansion, expressed by:

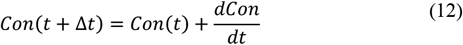

where *Con* is concentration of C, N, or O_2_ and 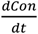 is the flux.

### Element fate calculation

We used Equation (13) to calculate element usage. Here, 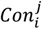 represents element *i* (includes C and N) usage concentration in *j* pathway (storage, respiration, biomass, N_2_ fixation). We calculated it by summing up the element usage in every time step, which is calculated by 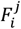 (the changing rate of element *i* in *j* pathway, e.g., C storage rate, C changing rate in respiration, C changing rate in growth) multiplied by the time step Δ*t*.

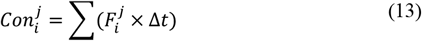

Equation (14) represents the calculation of the percentage of element usage. Here, 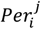 represents the percentage of *j* pathway (storage, respiration, biomass, N_2_ fixation) of element *i* (includes C and N) usage. 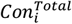 means the total element *i* usage.

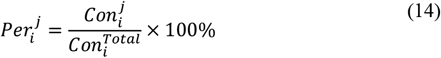

### Model simulation

In this study, we did four simulations. We simulated the model under H1 and H2, and under each hypothesis, we simulated two state switching conditions: a 1-minute switch and a 6-hour switch. All of these happened in 12 hours of daytime with light exposure.

### Parameters and tools

All parameters we used are included in the supplementary material (Table S1). Most required parameters are taken from a previous study ^3^, and we changed values in a reasonable range (Table S1).

## Supporting information

It includes two versions of the same supplementary. A pdf and a docx

It includes 2 videos, video 1 for H1 and video 2 for H2.

## Supplementary

Table S1: Parameters and values. Figure S1: O_2_ level and growth rate changes in 12 hours for H1. Figure S2: N_2_ fixation in 12 hours and 2 minutes.

## Code availability

https://doi.org/10.5281/zenodo.8062146

## Acknowledgements

M.G. and K.I. were supported by the U.S. National Science Foundation (OCE-2048373, subaward SUB0000525 from Princeton University). J. Z was supported by the Simons Foundation (awards 824082 and 72422 to J.P.Z.).

## Author contributions

Conceptualization: M.G., K.I., and J.Z.; Data curation: M.G.; Formal Analysis: M.G. and K.I.; Funding acquisition: K.I.; Investigation: M.G. and K.I.; Methodology: M.G., J.A. and K.I.; Project administration: K.I.; Resources: M.G., J.A., G.A., K.I., S.C. and J.Z.; Software: M.G.; Supervision: K.I.; Validation; Visualization: M.G., J.A, K.I. and S.C.; Writing – original draft: M.G., G.A. and K.I.; Writing – review & editing: M.G., G.A., K.I., S.C. and J.Z. All authors have read and agreed to the published version of the manuscript.

## Declaration of interests

The authors declare no competing interests.

## Notes

### Competing Interest Statement

The authors have declared no competing interest.

