## Supplementary material for "Rapid mode switching facilitates the growth of *Trichodesmium*: A model analysis": It includes two versions of the same supplementary. A pdf and a docx: Supplementary.docx

Table S1. Parameters and values.

| Parameter | Unit | Defination | Value |
| --- | --- | --- | --- |
| $C_{sto}$ | mol C mol C^-1^ | Carbon storage | Initial 0.5 |
| $F_{cfix}$ | d^-1^ | Carbon fixation rate |  |
| $F_{Bio}$ | d^-1^ | C changing rate in biomass (growth) |  |
| E | dimensionless unit | The ratio of respiration to biosynthesis | 0.4 |
| $F_{Cfix}^{max}$ | d^-1^ | Maximum carbon fixation rate |  |
| $A_{i}$ | µ mol^-1^ m^2^ s | Light saturation coefficient | 0.01 |
| *I* | µ mol m^-2^ s^-1^ | Light intensity | 700 |
| $F_{Bio}^{max}$ | d^-1^ | Maximum biomass production rate | 1 |
| $K_{C}$ | mol C mol C^-1^ | Half saturation constant of C storage | 0.2 |
| $K_{N}$ | mol N mol C^-1^ | Half saturation constant of N storage | 0.0318 |
| $\left[ O_{2} \right]$ | mol m^-3^ | Cellular oxygen concentration | Initial 0.213 |
| $\left[ O_{2} \right]_{E}$ | mol m^-3^ | Environmental oxygen concentration | 0.213 |
| $\rho_{C}^{Bio}$ | mol C m^-3^ | cellular C density | 18333 |
| $Y_{Cfix}^{O_{2}:C}$ | mol O_2_ mol C^-1^ | O_2_:C in photosynthesis | 1 |
| $N_{sto}$ | mol N mol C^-1^ | Nitrogen Storage | Initial 0.1 |
| A | d^-1^ | Diffusion coefficient of oxygen through cell membrane  layers | 311,040 |
| $\frac{dC_{sto}}{dt}$ | d^-1^ | C storage changing rate |  |
| $\frac{d\left[ O_{2} \right]}{dt}$ | mol m^-3^ d^-1^ | O_2_ concentration changing rate in *Trichodesmium* |  |
| $\frac{dN_{sto}}{dt}$ | d^-1^ | N storage changing rate |  |
| $Y_{Bio}^{N:C}$ | mol N mol C^-1^ | The ratio of N to C in biomass | 0.159 |
| $F_{N_{2}fix}$ | mol N mol C^-1^ d^-1^ | N_2_ fixation rate |  |
| $F_{N_{2}fix}^{max}$ | mol N mol C^-1^ d^-1^ | Maximum N_2_ fixation rate | 0.2 |
| $Y_{N_{2}fix}^{C:N}$ | mol C mol N^-1^ | The ratio of C to N in N_2_ fixation | 1 |
| $Y_{N_{2}fix}^{N:O_{2}}$ | mol N mol O_2_ ^-1^ | The ratio of N to O_2_ in N_2_ fixation | 1.29 |
| $Y_{Res}^{O_{2}:C}$ | mol O_2_ mol C ^-1^ | The ratio of O_2_ to C in N_2_ fixation | 1 |
| $F_{Res}$ | mol O_2_ m^-3^ d^-1^ | Respiration rate |  |
| $F_{Res}^{max}$ | mol O_2_ m^-3^ d^-1^ | Maximum respiration rate | 183,330 |
| $K_{O_{2}}$ | mol O_2_ m^-3^ | Half saturation concentration of O_2_ | 2×10^-5^ |

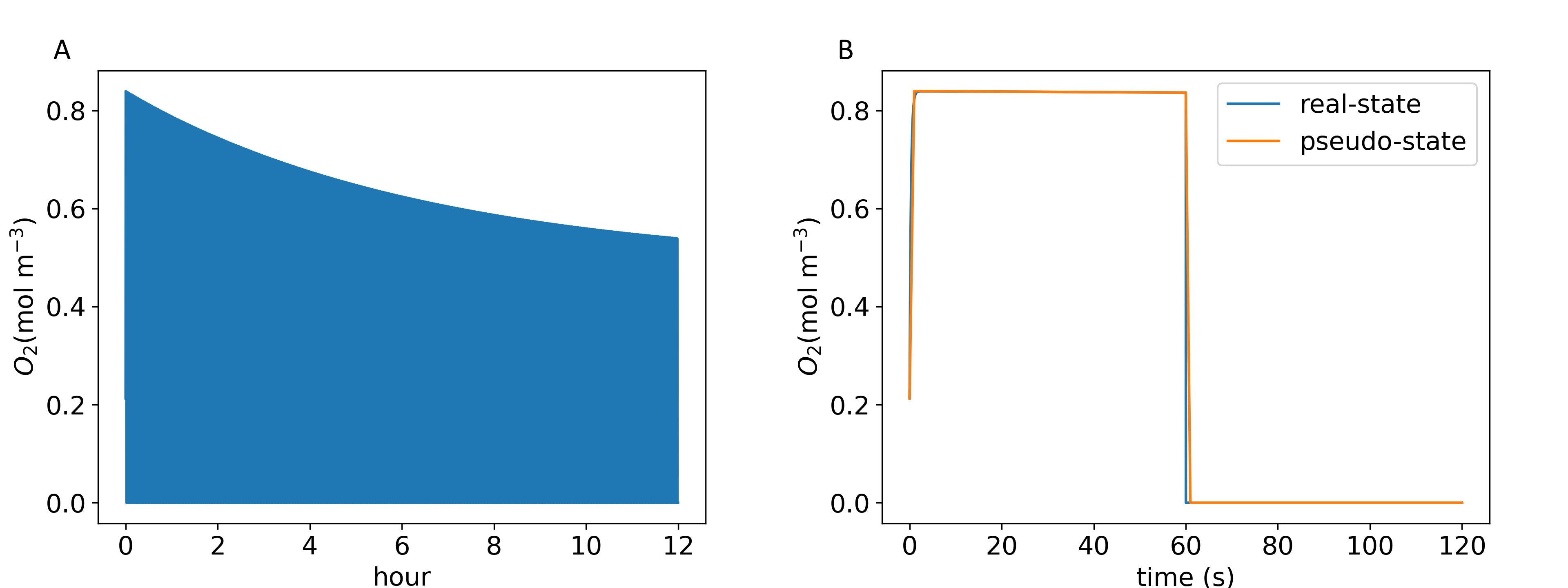

Figure S1. O_2_ level and growth rate changes in 12 hours for H1. A. Changes in O_2_ concentration in 12 hours. B. Changes in O_2_ concentration in 2 minutes; 60 s for photosynthetic state and 60 s for non-photosynthetic state.

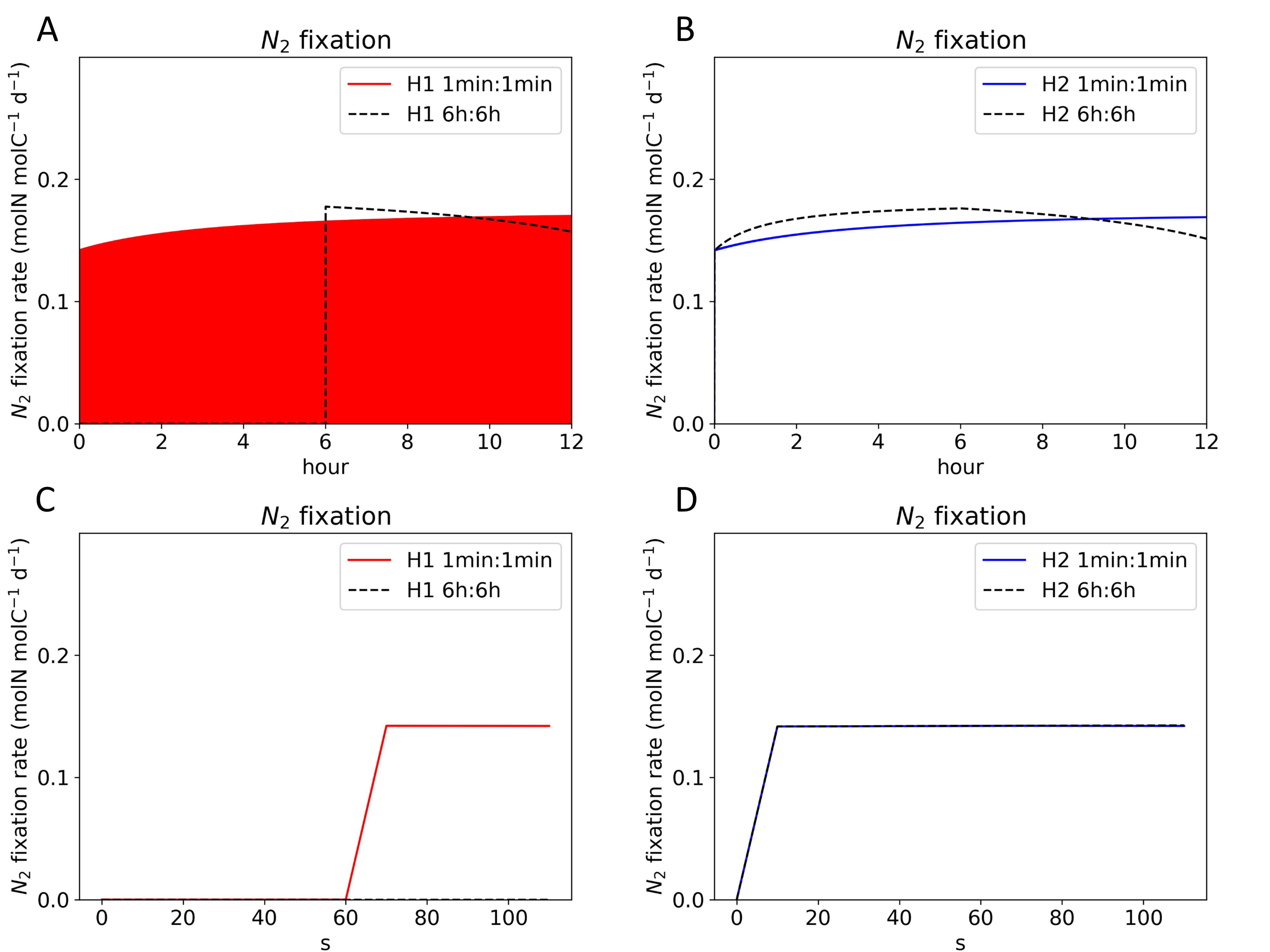

Figure S2. N_2_ fixation in 12 hours (A. H1 B. H2) and 2 minutes (C. H1 D. H2).
