## Supplementary material for "Rapid mode switching facilitates the growth of *Trichodesmium*: A model analysis": It includes two versions of the same supplementary. A pdf and a docx: Supplementary.pdf

Table S1. Parameters and values.

| Parameter | Unit | Defination | Value |
|---|---|---|---|
| $C_{sto}$ | mol C mol C$^{-1}$ | Carbon storage | Initial 0.5 |
| $F_{cfix}$ | d$^{-1}$ | Carbon fixation rate | |
| $F_{Bio}$ | d$^{-1}$ | C changing rate in biomass (growth) | |
| E | dimensionless unit | The ratio of respiration to biosynthesis | 0.4 |
| $F_{Cfix}^{max}$ | d$^{-1}$ | Maximum carbon fixation rate | |
| $A_i$ | μ mol$^{-1}$ m$^2$ s | Light saturation coefficient | 0.01 |
| $I$ | μ mol m$^{-2}$ s$^{-1}$ | Light intensity | 700 |
| $F_{Bio}^{max}$ | d$^{-1}$ | Maximum biomass production rate | 1 |
| $K_C$ | mol C mol C$^{-1}$ | Half saturation constant of C storage | 0.2 |
| $K_N$ | mol N mol C$^{-1}$ | Half saturation constant of N storage | 0.0318 |
| $[O_2]$ | mol m$^{-3}$ | Cellular oxygen concentration | Initial 0.213 |
| $[O_2]_E$ | mol m$^{-3}$ | Environmental oxygen concentration | 0.213 |
| $\rho_C^{Bio}$ | mol C m$^{-3}$ | cellular C density | 18333 |
| $Y_{Cfix}^{O_2:C}$ | mol $O_2$ mol C$^{-1}$ | $O_2$:C in photosynthesis | 1 |
| $N_{sto}$ | mol N mol C$^{-1}$ | Nitrogen Storage | Initial 0.1 |

| | | | |
|---|---|---|---|
| A | $d^{-1}$ | Diffusion coefficient of oxygen through cell membrane layers | 311,040 |
| $\frac{dC_{sto}}{dt}$ | $d^{-1}$ | C storage changing rate | |
| $\frac{d[O_2]}{dt}$ | mol $m^{-3}$ $d^{-1}$ | $O_2$ concentration changing rate in *Trichodesmium* | |
| $\frac{dN_{sto}}{dt}$ | $d^{-1}$ | N storage changing rate | |
| $Y_{Bio}^{N:C}$ | mol N mol $C^{-1}$ | The ratio of N to C in biomass | 0.159 |
| $F_{N_2fix}$ | mol N mol $C^{-1}$ $d^{-1}$ | $N_2$ fixation rate | |
| $F_{N_2fix}^{max}$ | mol N mol $C^{-1}$ $d^{-1}$ | Maximum $N_2$ fixation rate | 0.2 |
| $Y_{N_2fix}^{C:N}$ | mol C mol $N^{-1}$ | The ratio of C to N in $N_2$ fixation | 1 |
| $Y_{N_2fix}^{N:O_2}$ | mol N mol $O_2^{-1}$ | The ratio of N to $O_2$ in $N_2$ fixation | 1.29 |
| $Y_{Res}^{O_2:C}$ | mol $O_2$ mol $C^{-1}$ | The ratio of $O_2$ to C in $N_2$ fixation | 1 |
| $F_{Res}$ | mol $O_2$ $m^{-3}$ $d^{-1}$ | Respiration rate | |
| $F_{Res}^{max}$ | mol $O_2$ $m^{-3}$ $d^{-1}$ | Maximum respiration rate | 183,330 |
| $K_{O_2}$ | mol $O_2$ $m^{-3}$ | Half saturation concentration of $O_2$ | $2\times10^{-5}$ |

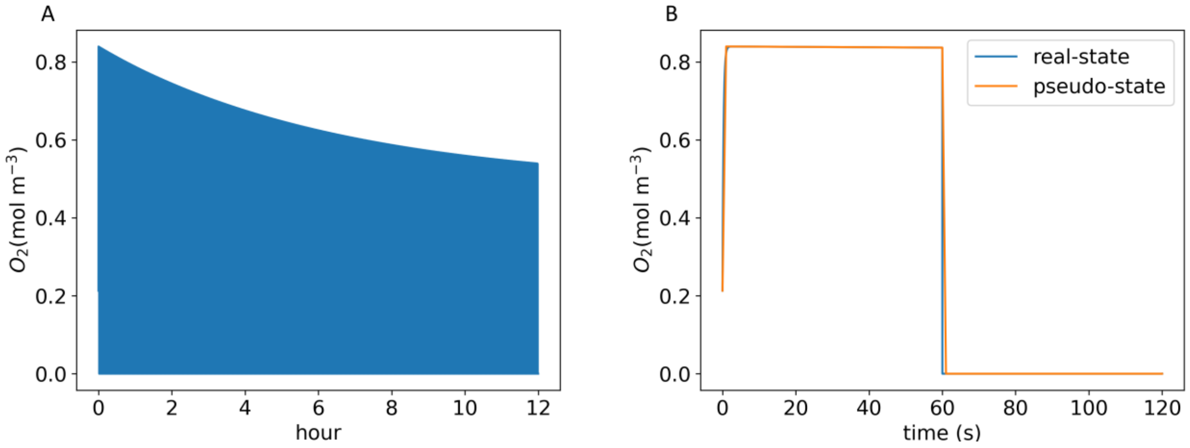


Figure S1. $O_2$ level and growth rate changes in 12 hours for H1. A. Changes in $O_2$ concentration in 12 hours. B. Changes in $O_2$ concentration in 2 minutes; 60 s for photosynthetic state and 60 s for non-photosynthetic state.

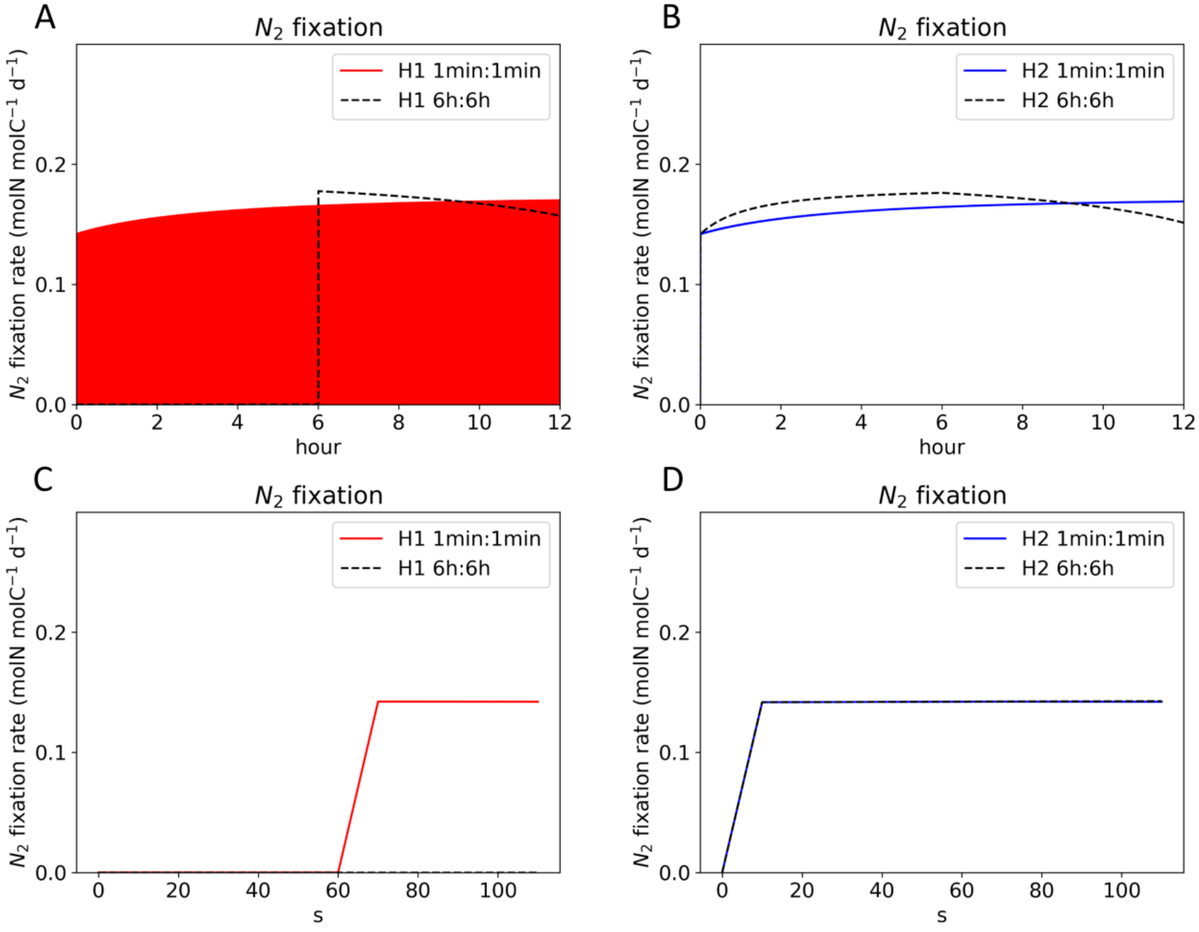


Figure S2. $N_2$ fixation in 12 hours (A. H1 B. H2) and 2 minutes (C. H1 D. H2).
